# TDP-43 pathology is sufficient to drive axon initial segment plasticity and hyperexcitability of spinal motoneurones in vivo in the TDP43-ΔNLS model of Amyotrophic Lateral Sclerosis

**DOI:** 10.1101/2024.06.01.596097

**Authors:** Svetlana Djukic, Zhenxiang Zhao, Lasse Mathias Holmsted Jørgensen, Anna Normann Bak, Dennis Bo Jensen, Claire Francesca Meehan

## Abstract

A hyperexcitability of the motor system is consistently observed in Amyotrophic Lateral Sclerosis (ALS) and has been implicated in the disease pathogenesis. What drives this hyperexcitability in the vast majority of patients is unknown. This is important to know as existing treatments simply reduce all neuronal excitability and fail to distinguish between pathological changes and important homeostatic changes. Understanding what drives the initial pathological changes could therefore provide better treatments. One challenge is that patients represent a heterogeneous population and the vast majority of cases are sporadic. One pathological feature that almost all (∼97%) cases (familial and sporadic) have in common is cytoplasmic aggregates of the protein TDP-43 which is normally located in the nucleus. In our experiments we investigated whether this pathology was sufficient to increase neuronal excitability and the mechanisms by which this occurs.

We used the TDP-43(ΔNLS) mouse model which successfully recapitulates this pathology in a controllable way. We used in vivo intracellular recordings in this model to demonstrate that TDP-43 pathology is sufficient to drive a severe hyper-excitability of spinal motoneurones. Reductions in soma size and a lengthening and constriction of axon initial segments were observed, which would contribute to enhanced excitability. Resuppression of the transgene resulted in a return to normal excitability parameters by 6-8 weeks. We therefore conclude that TDP-43 pathology itself is sufficient to drive a severe but reversible hyperexcitability of spinal motoneurones.

## Introduction

Despite decades of research, Amyotrophic Lateral Sclerosis (ALS) remains a fatal neurodegenerative disease with no cure. Until relatively recently, the only treatment strategy was a drug which reduces neuronal excitability (Riluzole). Although the effects of this drug are relatively limited, this heavily implicates excitotoxicity in the disease pathogenesis or progression. Transcranial magnetic stimulation, reflex and nerve excitability studies in individuals with ALS all confirm an increased excitability of the motor networks in this disease ^1–9^. One challenge with simply globally reducing neuronal excitability is that this cannot distinguish between pathological increases in neuronal excitability and homeostatic increases providing crucial compensation in a progressively weakening motor network. This potentially explains the limited effect of these drugs. An understanding of what is driving pathological excitability changes at a cellular level could likely produce more targeted and potentially more potent treatments. The proposed underlying mechanisms include an increased synaptic excitation and reduced inhibition of the motoneurones as well as an increased intrinsic excitability of the motoneurones, although what, in turn, drives these remains unknown. Our current experiments aimed to address just this.

A small proportion of cases are linked to mutations in the superoxide dismutase gene and mouse models based on ALS-associated SOD1 mutations have provided the main tool to date to investigate this topic *in vivo* with disease progression*. In vivo* intracellular recordings from spinal motoneurones in SOD1 mice have confirmed an increase in the intrinsic excitability of spinal motoneurones at a single cell level ^10–12^. Increased Na^+^ and Ca^2+^ persistent inward currents are seen in two different SOD1 models ^10,12–16^ which would both amplify synaptic input (contributing to the increased reflexes) and facilitate repetitive firing. Additionally, the current required to produce repetitive firing has been shown to be significantly reduced in SOD1 motoneurones with disease onset ^10,11,15^ and the gain of the motoneurones increased ^10,11^. Spontaneous action potential firing is also observed in these models ^10^.

Some of the proposed explanations for the increased excitability of motoneurones in SOD1 mice are decreases in soma size with disease progression in these mice ^17^, which could explain the increased input resistance also observed ^10^. Further, structural changes in the action potential generating region of the neurone - the axon initial segment, also appear to contribute. Early decreases in spinal motoneurone axon initial segment length seen in the G127X SOD1 mouse at the pre-symptomatic stage may reflect a homeostatic response to a more excitable motor cortex ^13^. As the disease progresses the axon initial segment increases in length and decreases in diameter in the same model ^12^, both of which would theoretically contribute to the increased motoneurone excitability seen in the disease.

However, SOD1 mutations account for a very small proportion (∼2%) of overall cases of ALS ^18^. What is driving the hyperexcitability in the vast majority of sporadic cases is therefore unclear. The most common key pathological feature found in both familial and sporadic cases involve the Tar-DNA binding protein TDP-43. The mis-location of TDP-43 from its normal location in the nucleus to the cytoplasm where it forms phosphorylated and highly ubiquitinated aggregates is the key pathological feature observed in around 97% of patients^19,20^. Recent work using neuronal cultured from induced pluripotent stem cells (IPSCs) from patients with TDP-43 mutations has demonstrated that these also show an increased neuronal excitability and axon initial segment changes^21^. Those experiments do potentially link TDP-43 with excitability changes in this disorder, although these cells represent a highly immature cell making it difficult to extrapolate to any adult disease stage.

Additionally, the excitability and axon initial segment length increases in the cells appear to precede actual TDP-43 pathology^21^. This is important to note as increased neuronal depolarization itself has been shown to drive TDP-43 cytoplasmic mis-location^22^. Whether similar changes in axon initial segments and spinal motoneurone excitability would therefore occur in the adult, in vivo, as a direct result of actual TDP-43 pathology is unknown.

To investigate this, we took advantage of the TDP-43(ΔNLS) mouse model which successfully recapitulates this pathology ^23^. This mouse contains a TDP-43 construct devoid of the nuclear localization signal, controlled by the Tet-Off system. This allows the transgene to be suppressed until adulthood then expressed resulting a relocation of TDP-43 from the nucleus to the cytoplasm leading to a severe and lethal ALS phenotype. This pathology and disease phenotype can be rescued upon subsequent transgene resuppression, allowing an *in vivo* “wash-out”. We used this model to investigate whether TDP-43 pathology is sufficient to induce hyper-excitability of spinal motoneurones *in vivo*. In these mice, we demonstrate that the mislocalisation and cytoplasmic aggregation of TDP-43 is sufficient to drive precisely the same excitability changes observed in familial mouse models of the disease explaining the hyperexcitability also seen in sporadic patients. Importantly, these changes were reversible upon transgene resuppression.

## Results

In the TDP-43 ΔNLS model a Tet-on/off system coupled to a TARDBP-gene missing the nuclear localisation signal allowed a controllable expression of this transgene (**Fig. 1A**). Crossing this with mice expressing the tTA transgene downstream of the Neurofilament Heavy Chain (NEFH) promoter, the controllable expression was obtained in all neurons. We used doxycycline food to suppress the transgene until the mice reached adulthood. Removal of the doxycycline diet at 7 weeks of age then resulted in a loss of nuclear TDP-43, and cytoplasmic accumulation of phosphorylated TDP-43 (**Fig.1B**) as has been previously reported for this model ^23^. This resulted in a severe and progressive motor phenotype.

**Figure 1.**
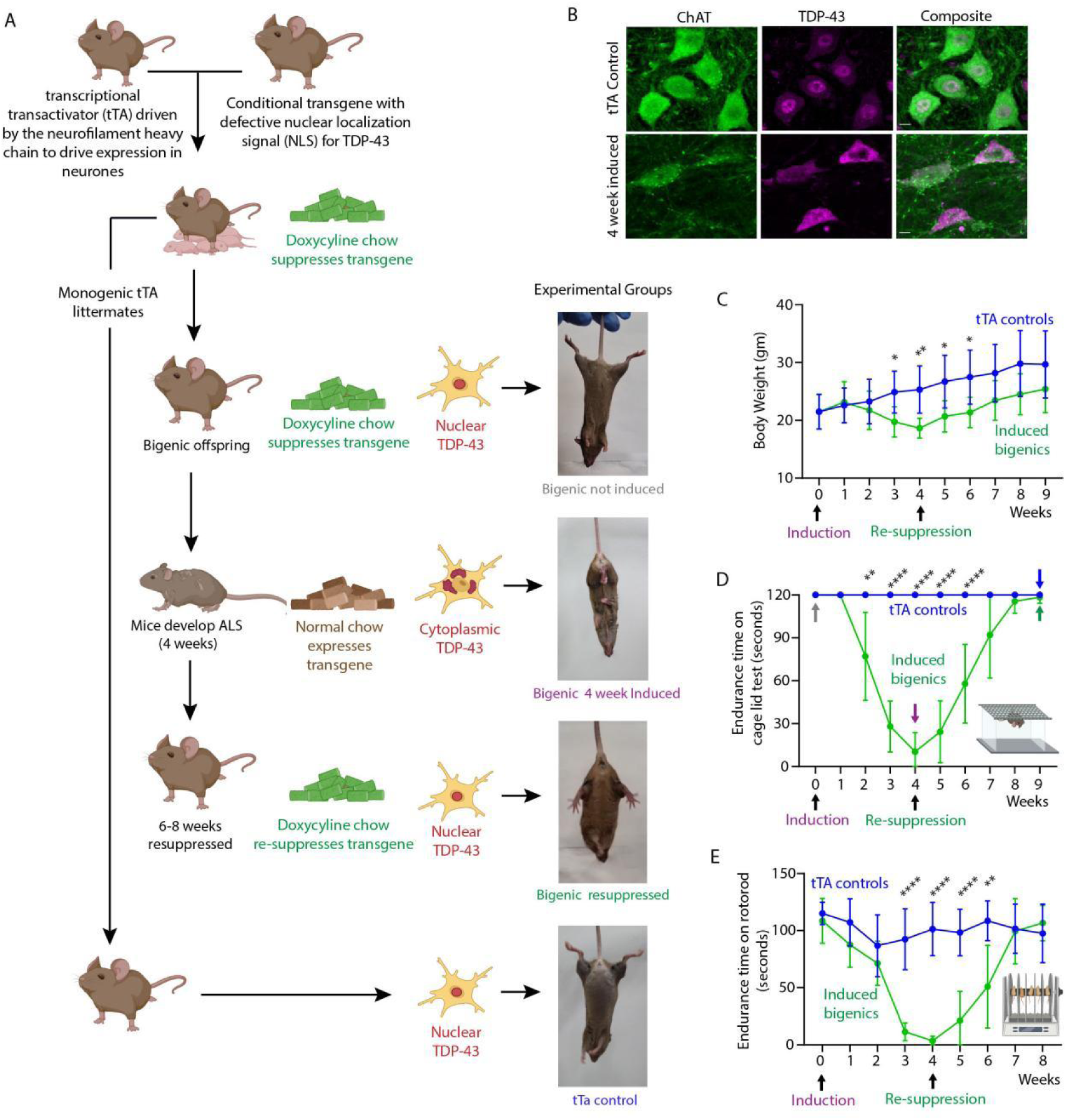
TDP-43 Pathology drives a reversible ALS phenotype. (**A**) Breeding schema and experimental groups including a bigenic not-induced control group, a bigenic 4 weeks induced group, a bigenic resuppressed group, and a tTA control group. Representative pictures from the tail suspension test are shown demonstrating the reversible ALS phenotype seen in induced bigenic mice. By 4 weeks post-induction mice showed a severe phenotype with extreme clasping of all 4 limbs, and recovered after 6-8 weeks of resuppression. (**B)** Representative confocal images of motoneurones (labelled with antibodies against ChAT, green) showing a loss of nuclear TDP-43, and cytoplasmic accumulation of phosphorylated TDP-43 (magenta), after removing of the doxycycline diet at 7 weeks of age (scale bar is 10 µm). (**C**) Bigenic mice (green) show extreme weight loss after induction and recovered after resuppression, compared to tTA control littermates. Mixed –effect analysis Šídák’s multiple comparisons test, P= 0.0283 (3 weeks), P= 0.0096 (4 weeks), P=0.0411 (1 week after resuppression) and P=0.0425 (2 weeks after resuppression). N=9 tTA mice (5 female, 4 male) and 12 bigenic mice (4 female, 8 male). (**D**) Endurance time on cage grid test shows that bigenic mice (green) decrease greatly after induction and recover after resuppression, while the tTA group (blue) shows no change. Mixed –effect analysis Šídák’s multiple comparisons test, P= 0.0052 (2 weeks), P<0.0001 (3 and 4 weeks post induction and 1 and 2 weeks after resuppression). N=9 tTA mice (5 female, 4 male) and 12 bigenic mice (4 female, 8 male). (**E**) Endurance time on rotarod shows that bigenic mice (green) decrease greatly after induction and recovered after resuppression, while the tTA group (blue) shows no changes. Mixed –effect analysis Šídák’s multiple comparisons test, P= 0.0052 (2 weeks), P<0.0001 (3 and 4 weeks post induction and 1 week after resuppression) and P=0.0015 (2 weeks after resuppression). N=9 tTA mice (5 female, 4 male) and 12 bigenic mice (4 female, 8 male). Data are given as the mean ±SD. \**P* < 0.05, \*\**P* < 0.01, \*\*\**P* < 0.001, \*\*\*\**P* < 0.0001

By two weeks post-induction mice began to clasp in their hind-limbs when suspended by the tail (**Fig. 1A**) and already showed significantly reduced compound muscle action potentials compared to monogenic controls. By 4 weeks mice showed an even more extreme phenotype with extreme clasping of all 4 limbs and extreme weight loss (**Fig.1C**), muscle atrophy and weakness. This could be quantified by the behavioural tests including the endurance time that the mice could hang on to a steel grid placed 30cm above a padded surface (**Fig. 1D**) and on the rotarod task (**Fig. 1E**). Bigenic mice maintained on the doxycyline diet (“not-induced mice”) did not show this change. Neither did monogenic tTA only mice of the same age that went through the same feeding regime as the expressed mice (tTA only controls). Males showed a faster disease progression than females on the cage grid task (**Suppl. Fig. 1A**) and some were even close to humane endpoint by 4 weeks. Terminal experiments were therefore performed on one group of bigenic mice at 4 weeks post-induction (11 weeks of age) when the mice showed this severe motor phenotype. In a second cohort of bigenic mice after 4 weeks of transgene induction the transgene was then resuppressed by the reintroduction of doxycycline to the diet. This resulted in a significant recovery of motor function by 6 weeks after resuppression. Although resuppressed mice still showed a relatively mild clasping phenotype, they performed equally well as controls on both the grid endurance task (**Fig. 1D**) and the rotarod task (**Fig. 1E**) along with regaining their weight (**Fig.1C**). Therefore, experiments were also performed at 6 weeks post resuppression. Two controls groups were used consisting of age-matched (10 week old) bigenic mice with a suppressed transgene (on the doxycycline food) and 17 week old monogenic tTA only littermate mice that had gone through the same doxycycline regime (doxycycline removal for 4 weeks and reinstatement for 6 weeks). This latter group also controlled for any potential toxicity of the expression of the tTA transactivator.

### TDP-43 pathology drives a pathological motoneurone hyperexcitability in vivo

To measure the effect of the TDP-43 pathology on neuronal excitability, intracellular recording was performed on these 4 groups. Here, a microelectrode was inserted into antidromically identified spinal motoneurones in vivo in anaesthetized mice (**Fig. 2A**). Once inside the cell, current was injected through the microelectrode into the cell to mimic synaptic input and the neuronal response was recorded. The current strength was increased in a linear fashion using triangular ramps and the current level required to evoke repetitive firing (the rheobase) was measured for each cell (examples shown in **Fig. 2B and C**). The rheobase was significantly lower after 4 weeks of transgene induction than in bigenic not-induced controls by approximately 74% (**Fig. 2D**). Resuppression of the transgene resulted in a complete restoration of the control rheobase levels and tTA only controls confirmed that the decrease in rheobase was not a consequence of the tTA expression alone.

**Figure 2.**
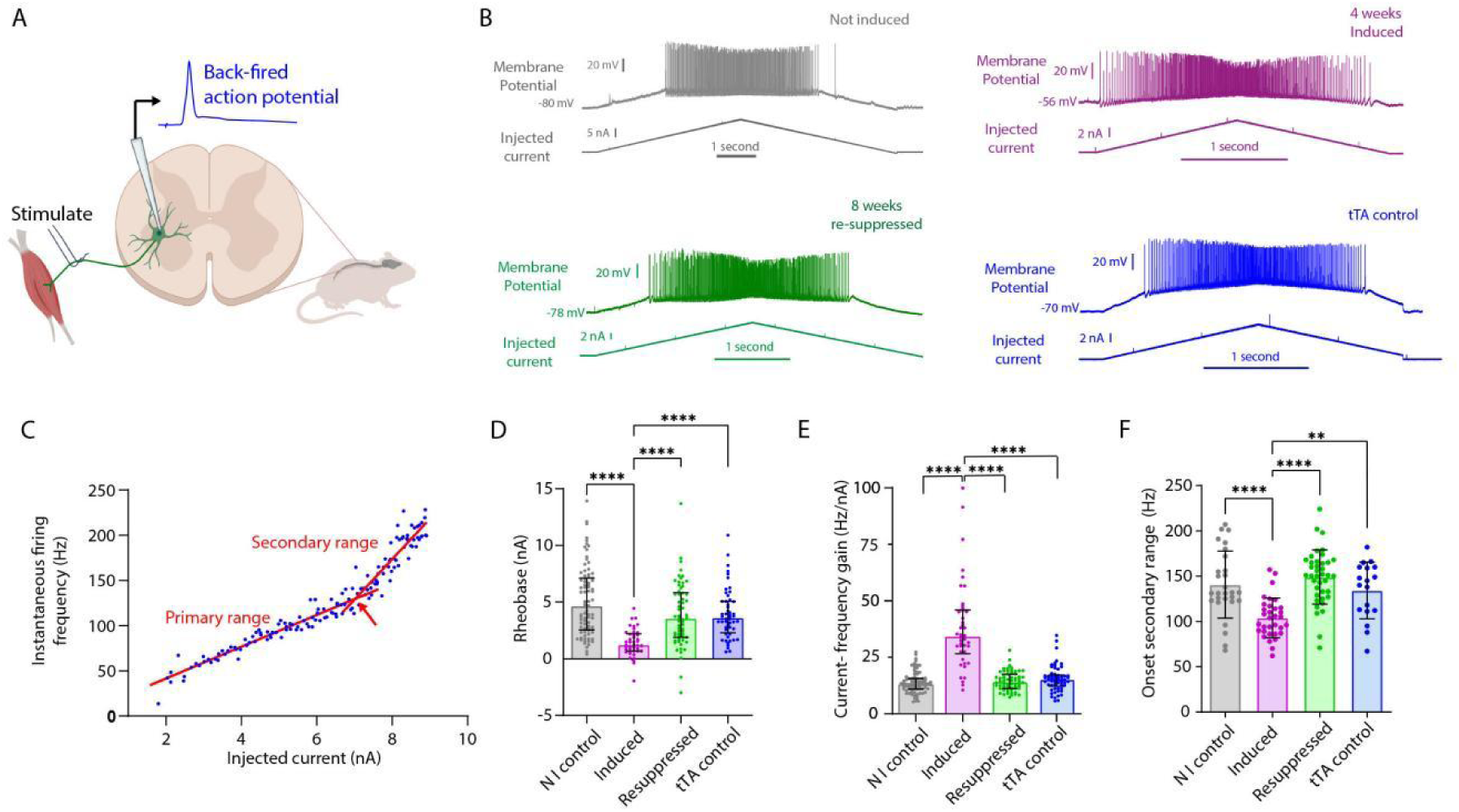
TDP-43 pathology drives a reversible hyperexcitability of spinal motoneurones. (**A**) Cartoon of the experimental setup for the *in vivo* intracellular recording. Stimulation of the distal peripheral nerve initiates antidromic action potentials in motor axons towards the spinal cord. The antidromic action potentials in motoneurones (blue trace) are recorded intracellularly. (**B**) Representative voltage traces (see membrane potential) showing the spike discharge frequency in response to triangular intracellular injections of current in motoneurones. Examples are shown for each of the four different groups (Bigenic not-induced (grey), Bigenic induced (magenta), Bigenic resuppressed (green) and tTA control (blue)). (**C**) Representative current-frequency gain to illustrate the primary range and secondary range. Red arrow indicates the transition between the two ranges. (**D**) Scatter dot plot showing the recruitment currents (rheobase) for the four different groups displayed. This shows the rheobase decreased significantly in the induced mice and returned to control values upon resuppression. Plot shows medians and interquartile ranges and each dot shows the rheobase of a single neurone. Medians (and IQR), Not-induced: 4.635nA (4.584), Induced: 1.215nA (1.543), Resuppressed: 3.54nA (5.838), tTA control: 3.6nA (2.773). Kruskal Wallis, P <0.0001, Not-induced: n = 82 cells (6 mice, 3 female, 3 male), Induced: n= 42 (9 mice, 5 female, 4 male), Resuppressed: n= 62 cells (6 mice 3 female, 3 male) and tTA control: n=52 cells (5 mice, 3 female, 2 male). Dunn’s Multiple comparisons tests: Not-induced vs. Induced P <0.0001, Induced vs. resuppressed P <0.0001, Induced vs tTA P <0.0001. Other pairwise comparisons were not significant. (**E**) Scatter dot plot showing the current-frequency (I-f) gains measured in the primary range from the four different groups. This increased significantly in the induced mice and returned to control values upon resuppression. Plot shows medians and interquartile ranges and each dot shows the primary range I-f slope of a single neurone in the data set. Medians (and IQR), Not-induced: 13.12 Hz/nA (4.65), Induced: 34.22 Hz/nA (19.35), Resuppressed: 14.05 Hz/nA (6.30), tTA control: 15.12 Hz/nA (4.97). Kruskal Wallis, P <0.0001, Not-induced: n = 82 cells (6 mice, 3 female, 3 male), Induced: n= 42 (9 mice, 5 female, 4 male), Resuppressed: n= 62 cells (6 mice, 3 female, 3 male) and tTA control: n=58 cells (5 mice, 3 female, 2 male). Dunn’s Multiple comparisons tests: Not-induced vs. Induced P <0.0001, Induced vs. resuppressed P <0.0001, Induced vs tTA P <0.0001. Other pairwise comparisons were not significant. (**F**) Scatter dot plot showing the onset of the secondary range for the four different groups. This decreased significantly in the induced mice and returned to control values upon resuppression. Plot shows means and standard deviations and each dot shows the firing frequency at the onset of the secondary of a single neurone in the data set. Means (and SDs), Not-induced: 140.06 Hz (36.93), Induced: 104.00 Hz (21.91), Resuppressed: 149.00 Hz (29.99), tTA control: 134.20 Hz (31.34). Kruskal Wallis, P <0.0001, Not-induced: n = 28 cells (6 mice, 3 female, 3 male), Induced: n= 33 cells (8 mice, 5 female, 3 male), Resuppressed: n= 39 cells (6 mice, 3 female, 3 male) and tTA control: n=18 cells (5 mice, 3 female, 2 male). Tukey’s multiple comparisons tests: Not-induced vs. Induced P <0.0001, Induced vs. resuppressed P <0.0001, Induced vs tTA P <0.0045. Other pairwise comparisons were not significant. *P < 0.05, **P < 0.01, ***P < 0.001, ****P < 0.0001

As the current level increased, the instantaneous firing frequency was calculated and the gain of the motoneurones obtained. This slope had two phases; the primary range and the secondary range (**Fig. 2C**). The gain of the motoneurone was calculated in the primary range and this was significantly higher in the 4-week induced group compared to the not-induced group by 160%. Once again, this value returned to baseline levels subsequent to transgene resuppression and was also not significantly different in tTA controls (**Fig. 2E**). The secondary range represents a sharp inflection in the current-frequency (I-f) slope. This is believed to reflect the activation of dendritic persistent inward currents, which amplify the inputs, thereby increasing the I-f gain. The onset of these has been shown to be at significantly lower firing frequencies in SOD1 models of the disease ^10,12,13,16^ and so this was also measured. As seen in SOD1 models, the onset of the secondary range occurred at a significantly lower firing frequency in the induced mice compared to both the not-induced bigenic mice and the tTA controls. Again, this value returned to normal upon resuppression (**Fig 2F**). Thus, we can conclude that the observed changes mentioned above result in an intrinsically more excitable motoneurone. This was observed regardless of the sex of the mice (**Supp. Fig. 1B** and **C**).

### Reductions in soma size and increased input resistance contribute to the increased excitability

To investigate anatomical features that could explain the increased intrinsic excitability, we first analysed the soma size of retrogradely traced soleus and gastrocnemius motoneurones (**Fig. 3A**). Here, a significant reduction in soma size was observed for gastrocnemius motoneurones in induced mice compared to both not-induced bigenic mice and the tTA controls. Once again, this returned to control (tTA only) values upon resuppression (**Fig. 3B**). Although here it must be noted that the bigenic not-induced gastrocnemius motoneurones had a larger average soma size than the tTA controls suggesting that either the tTA or/and doxycycline treatment may induce changes in soma size. For soleus motoneurones, a significant decrease in soma size was also seen in induced mice relative to the bigenic not-induced mice (**Fig. 3C**). Although, this was not significantly different from the tTA controls, suggesting that the reduction in soma size is not as extreme at this time point in soleus motoneurones. Reductions in soma size could, in theory lead to increased input resistance and so this was tested electrophysiologically using -3nA hyperpolarizing pulses (**Fig. 3D**). This confirmed a large significant increase in input resistance in induced mice compared to both not-induced bigenic mice and tTA controls. Once again, this returned to normal values upon resuppression (**Fig. 3E**). Thus reductions in soma size drive increases in input resistance which could help to explain both the decreased rheobase and the increased gain seen in the induced mice. A side effect of this increased input resistance was an increased activation of hyperpolarization-activated currents (I_h_ currents), measured indirectly as the sag seen in the voltage response towards the end of the current pulse (**Fig. 3F**). Although this was not significantly different when expressed as a proportion of the overall voltage drop, the larger voltage drop in the induced mice resulted in a larger absolute sag.

**Figure 3.**
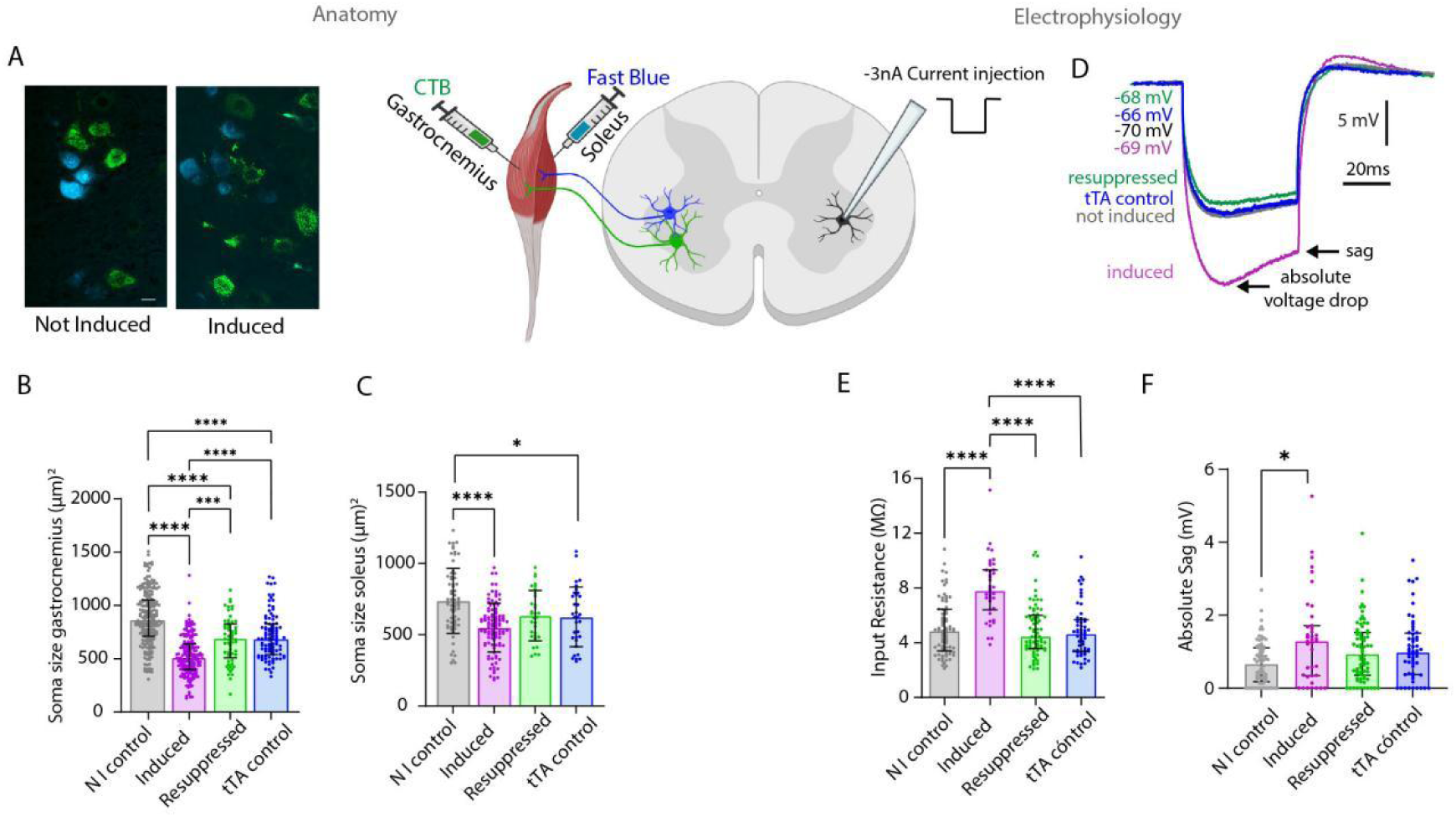
Soma size contributes to an increased input resistance in induced mice. (**A**) Representative confocal images of gastrocnemius motoneurones (green) and soleus motoneurones (cyan) retrogradely traced using the Cholera Toxin β subunit conjugated to Alexa 488 (CTβ-488) and Fast Blue, respectively. (**B**) Scatter dot plots showing the 2D Soma size measurements for gastrocnemius motoneurones from the four different groups. This shows that the soma size decreased significantly in the induced mice compared to both the not-induced bigenic mice and tTa only controls, but returned to control (tTA only) values upon resuppression. Plot shows medians and interquartile ranges and each dot shows a single neurone. Medians (and IQR), Not-induced: 861.4 μm^2^(340.2), Induced: 507.6 μm^2^ (239.6), Resuppressed: 687.7 μm^2^ (317.6), tTA control: 687.1 μm^2^ (288,2). Kruskal Wallis, P <0.0001, Not-induced: n = 208 cells (8 mice, 4 female 4 male), Induced: n= 181 (8 mice, 5 female, 3 male), Resuppressed: n= 57 cells (4 mice, 2 female, 2 male) and tTA control: n=95 cells (6 mice, 3 female, 3 male). Dunn’s Multiple comparisons tests: Not-induced vs. Induced P <0.0001, Not-induced vs. resuppressed P <0.0001, Induced vs. resuppressed P <0.001, Not-induced vs tTA P <0.0001, induced vs tTA P <0.0001. Other pairwise comparisons were not significant. (**C**) Scatter dot plots showing the 2D Soma size measurements for soleus from the four different groups. This shows that the soma size for these cells decreased significantly in the induced mice compared to both the not-induced bigenic mice and tTA only controls, but returned to control (tTA only) values upon resuppression. Plot shows means and standard deviations and each dot shows a single neurone. Mean (and SD), Not-induced: 738.2 μm^2^ (227.9), Induced: 549.9 μm^2^ (171.5), Resuppressed: 634.3 μm^2^ (176.6), tTA control: 625.2 μm^2^ (209.5). ANOVA, P <0.0001, Not-induced: n = 67 cells (8 mice, 4 female, 4 male), Induced: n= 102 (7 mice, 4 female, 3 male), Resuppressed: n= 32 cells (6 mice, 3 female, 3 male) and tTA control: n=33 cells (4 mice, 3 female, 1 male). Tukey’s Multiple comparisons tests: Not-induced vs. Induced P <0.0001, Not-induced vs tTA P <0.05. Other pairwise comparisons were not significant. (**D**) Representative examples of the voltage drop in response to -3nA hyperpolarising current pulses to test input resistance. (**E**) Scatter dot plot showing input resistance from the four different groups. The input resistance increased significantly in the induced mice compared to both the not-induced bigenic mice and the tTA only controls, and returned to both control values upon resuppression. Plot shows medians and interquartile ranges and each dot shows the input resistance of a single neurone. Kruskal Wallis, P <0.0001. Medians (and IQR), Not-induced: 4.85 MΩ (3.04), Induced: 7.78 MΩ (2.92), Resuppressed: 4.45 MΩ (2.42), tTA control: 4.63 MΩ (2.29). Kruskal Wallis, P <0.0001, Not-induced: n = 74 cells (6 mice, 3 female, 3 male), Induced: n= 39 (9 mice, 5 female, 4 male), Resuppressed: n= 72 cells (6 mice, 3 female, 3 male) and tTA control: n= 56 cells (5 mice, 3 female, 2 male). Dunn’s Multiple comparisons tests: Not-induced vs. Induced P <0.0001, Induced vs. resuppressed P <0.0001, Induced vs tTA P <0.0001. Other pairwise comparisons were not significant. (**F**) Scatter dot plot showing the absolute sag for the four different groups. This suggests that the increased voltage drop in induced mice activate more Ih current. Plot shows medians and interquartile ranges and each dot shows the input resistance of a single neurone. Kruskal Wallis, P=0.0088. Medians (and IQR), Not-induced: 0.66mV (0.83), Induced: 1.29 mV (1.37), Resuppressed: 0.94 mV (1.17), tTA control: 0.98 mV (1.13). Kruskal Wallis, P <0.0001, Not-induced: n = 74 cells (6 mice, 3 female, 3 male), Induced: n= 39 (9 mice, 5 female, 4 male), Resuppressed: n= 72 cells (6 mice, 3 female, 3 male) and tTA control: n= 56 cells (5 mice, 3 female, 2 male). Dunn’s Multiple comparisons tests: Not-induced vs. Induced P <0.05. Other pairwise comparisons were not significant. *P < 0.05, **P < 0.01, ***P < 0.001, ****P < 0.0001

### Axon initial segment changes are also consistent with an increased motoneurone excitability

We next investigated axon initial segment size and location in traced motoneurones (**Fig. 4**) as this has also been shown to influence repetitive firing ^24^. Axon initial segments were immunohistochemically labelled using antibodies against Ankyrin G, the main scaffolding protein that spans the entire axon initial segment (**Fig. 4**). As fast motoneurones are known to be more vulnerable in this disease, we injected different tracers into the predominately slow twitch soleus muscles (fast blue tracer) and the predominately more fast twitch gastrocnemius muscles (CTB tracer). Importantly, by injecting the tracer into the muscle, this ensures that the motoneurones were still functionally connected to their respective muscle as a functional block at the NMJ can also affect axon initial segment length ^25^. First, we noticed that many cells (in particular, the gastrocnemius motoneurones) had swollen axon hillocks where the tracer aggregated and often failed to make it all the way into the soma (**Fig. 4E**).

**Figure 4.**
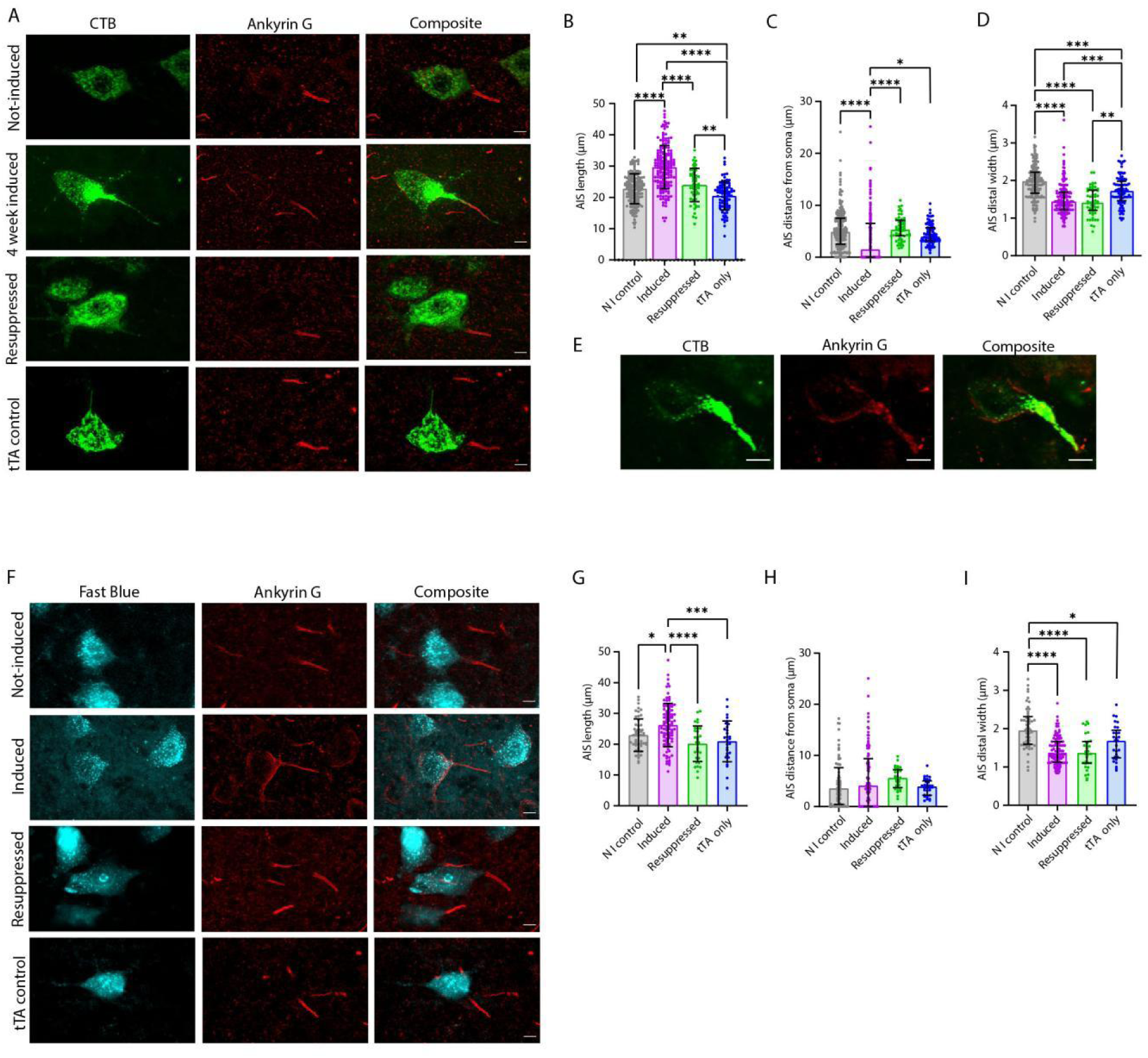
TDP-43 pathology drives reversible increases in axon initial segment length on both gastrocnemius and soleus motoneurones. (**A**) Representative confocal images of gastrocnemius motoneurones from the four groups, identified by retrograde tracing using the Cholera Toxin β subunit conjugated to Alexa 488 (CTβ-488, in green). Axon initial segments were immunohistochemically labelled using antibodies against Ankyrin G (red). Scale bar indicates 10 µm. (**B**) Scatter dot plot showing an increase in axon initial segment length on gastrocnemius motoneurones after transgene induction that returns to control values after resuppression. Plot shows means and standard deviations and each dot shows a single neurone. Mean (and SD), Not-induced: 22.76 µm (4.79), Induced: 29.67 µm (6.90), Resuppressed: 23.98 µm (5.31), tTA control: 21.2 µm (4.49). ANOVA, P <0.0001, Not-induced: n = 179 cells (8 mice, 5 female, 3 male), Induced: n= 181 (6 mice, 3 female, 3 male), Resuppressed: n= 57 cells (5 mice, 2 female, 3 male,) and tTA control: n=94 cells (6 mice, 3 female, 3 male). Tukey’s Multiple comparisons tests: Not-induced vs. Induced P <0.0001, Induced vs. resuppressed P <0.001, Not-induced vs tTA P <0.01. Induced vs tTA P <0.001 resuppressed vs tTA control P <0.01. Other pairwise comparisons were not significant. (**C**) Scatter dot plot showing a decrease in the distance of the proximal axon initial segment from the soma of gastrocnemius motoneurones after transgene induction. This also returns to control value upon resuppression. Plot shows medians and interquartile ranges and each dot shows a single neurone. Medians (and IQR), Not-induced: 4.92 µm (34.99), Induced: 1.56 µm (6.51), Resuppressed: 5.36 µm (3.01), tTA control: 3.98 µm (2.51). Kruskal Wallis, P <0.0001, Not-induced: n = 212 cells (8 mice, 5 female, 3 male), Induced: n= 181 (6 mice, 3 female, 3 male), Resuppressed: n= 56 cells (5 mice, 2 female, 3 male) and tTA control: n=95 cells (6 mice, 3 female, 3 male). Dunn’s Multiple comparisons tests: Not-induced vs. Induced P <0.0001, Induced vs. resuppressed P <0.001, Induced vs tTA P <0.05. Other pairwise comparisons were not significant. (**D**) Scatter dot plot showing that distal diameter of the axon initial segment on gastrocnemius motoneurones is reduced after transgene induction. This does not return to normal value upon resuppression. Plot shows medians and interquartile ranges and each dot shows a single neurone. Medians (and IQR), Not-induced: 1.98 µm (1.31), Induced: 1.44 µm (0.91), Resuppressed: 1.21 µm (1.1), tTA control: 1.72 µm (1.03). Kruskal Wallis, P <0.0001, Not-induced: n = 179 cells (8 mice, 5 female, 3 male), Induced: n= 181 (6 mice, 3 female, 3 male), Resuppressed: n= 57 cells (5 mice, 2 female, 3 male) and tTA control: n=95 cells (6 mice, 3 female, 3 male). Dunn’s Multiple comparisons tests: Not-induced vs. Induced P <0.0001, Induced vs. resuppressed P <0.0001, Not-induced vs tTA control P <0.001, Induced vs tTA control P <0.001, resuppressed vs tTA control P <0.01. Other pairwise comparisons were not significant. (**E**) Confocal images (maximum projection) of a traced gastrocnemius motoneurone, showing an accumulation of the tracer at the axon hillock and a breakdown of the proximal border of the axon initial segment with the Ankyrin G extending into the soma. Scale bar indicates 10 µm. (**F**) Representative confocal images (maximum projections) of soleus motoneurones from the four groups, identified by retrograde tracing using Fast Blue tracer (in cyan). A breakdown of a proximal border can be seen in the induced example here. Scale bar indicates 10 µm. (**G**) Scatter dot plot showing an increase in axon initial segment length also occurs in soleus motoneurones after transgene induction and returns to control values after resuppression. Although, this is less extreme than seen in gastrocnemius motoneurones. Plot shows means and standard deviations and each dot shows a single neurone. Mean (and SD), Not-induced: 22.95 µm (5.28), Induced: 26.22 µm (7.07), Resuppressed: 20.13 µm (5.76), tTA control: 20.93 µm (6.64). ANOVA, P <0.0001, Not-induced: n = 53 cells (8 mice, 5 female 3 male), Induced: n= 102 (6 mice, 3 female, 3 male), Resuppressed: n= 32 cells (56 mice, 3 female, 3 male,) and tTA control: n= 30 cells (6 mice, 3 female, 3 male). Tukey’s Multiple comparisons tests: Not-induced vs. Induced P=0.0157, Induced vs. resuppressed P <0.0001, Induced vs tTA control P=0.0006. Other pairwise comparisons were not significant. (**H**) Scatter dot plot showing no changes in the distance of the proximal axon initial segment from the soma in soleus motoneurones after transgene induction. Plot shows medians and interquartile ranges and each dot shows a single neurone. Medians (and IQR), Not-induced: 3.55 µm (7.18), Induced: 4.08 µm (9.38), Resuppressed: 5.58 µm (3.52), tTA control: 3.91 µm (2.79). Kruskal Wallis, P <0.1334, Not-induced: n = 66 cells (8 mice, 5 female 3 male), Induced: n= 102 (6 mice, 3 female, 3 male), Resuppressed: n= 32 cells (6 mice, 3 female, 3 male) and tTA control: n=33 cells (6 mice, 3 female, 3 male). (**I**) Scatter dot plot showing that distal width of the axon initial segment on gastrocnemius motoneurones is reduced after transgene inductions. This does not return to normal value upon resuppression. Plot shows medians and interquartile ranges and each dot shows a single neurone. Medians (and IQR), Not-induced: 1.96 µm (0.72), Induced: 1.34 µm (0.54), Resuppressed: 1.37 µm (0.57), tTA control: 1.24 µm (0.72). Kruskal Wallis, P <0.0001, Not-induced: n = 53 cells (8 mice, 5 female 3 male), Induced: n= 102 (6 mice, 3 female, 3 male), Resuppressed: n= 32 cells (6 mice, 3 female, 3 male) and tTA control: n= 29 cells (6 mice, 3 female, 3 male). Dunn’s Multiple comparisons tests: Not-induced vs. Induced P <0.0001, Not-induced vs Resuppressed P <0.0001, Not-Induced vs tTA control P =0.0229. Other pairwise comparisons were not significant.

Starting with the most vulnerable gastrocnemius motoneurones (**Fig. 4A**), our results confirmed that axon initial segments were significantly longer in the induced mice compared to both the not-induced bigenic mice and the tTA only controls (**Fig. 4B**). This appeared to be due to a breakdown of the proximal border of the axon initial segment and the axon hillock, which was seen as the Ankyrin G labelling extending into the soma in some cases (**Fig. 4E**). Not surprisingly, on average, the axon initial segment was significantly closer to the soma on the induced gastrocnemius (**Fig 4C**). However, the distal axon initial segment border was also significantly further away confirming that the axon initial segments elongated both distally and proximally in the induced mice. Both of these parameters returned to normal values upon resuppression. Another key feature that was observed was that, despite the proximal axon swellings, the distal end of axon initial segments had a significantly smaller diameter in the induced mice compared to both the not-induced bigenic mice and the tTA only controls. However, this parameter did not return to control value upon resuppression (**Fig. 4D**).

Next, axon initial segments on the less vulnerable soleus motoneurones were measured (**Fig. 4F**). These were also significantly longer in induced mice and returned to control values following transgene resuppression (**Fig. 4G**). For these the distance from the soma was not significantly different, suggesting that the elongation mostly occurred in the distal end (**Fig. 4H**), although in a few cases, a breakdown of the proximal border was also seen for soleus motoneurones (**Fig. 4F**) however, these were not as common as for gastrocnemius. A permanent reduction in axon initial segment width was also observed (**Fig. 4I**).

Together, the reduction in soma size, increase in input resistance and elongation and constriction of the axon initial segments could be predicted to result in an increased excitability of the motoneurones. This is consistent with our electrophysiological results.

### TDP-43 pathology alters individual action potentials

Finally, we turned our focus to the features of individual actions potentials to determine the effect of the axon initial segment changes on these. For this, we first analysed the antidromic action potential, examples of which are shown in **Fig. 5A**. Surprisingly, the amplitudes of the somatic portion of the spike were significantly lower in the induced mice than both not-induced bigenic mice and tTA controls, but returned to normal values upon resuppression (**Fig. 5B**). The antidromic spikes were then differentiated to reveal the maximal rate of rise of the spike as it arrives at the axon initial segment (IS spike) versus the soma and dendrites (the SD spike) (**Fig 5C**). Consistent with the reduced amplitude of the somatic spike, the maximum rates of rise of the SD spike were significantly lower in the induced mice than both controls but, once again, returned to normal values upon resuppression (**Fig. 5D**). This is consistent with decreased fast transient Na+ conductance in the soma. This was not observed for the IS spike (**Fig. 5E**) suggesting and altered balance between Na+ channels in the soma and the axon initial segment in induced mice.

**Figure 5.**
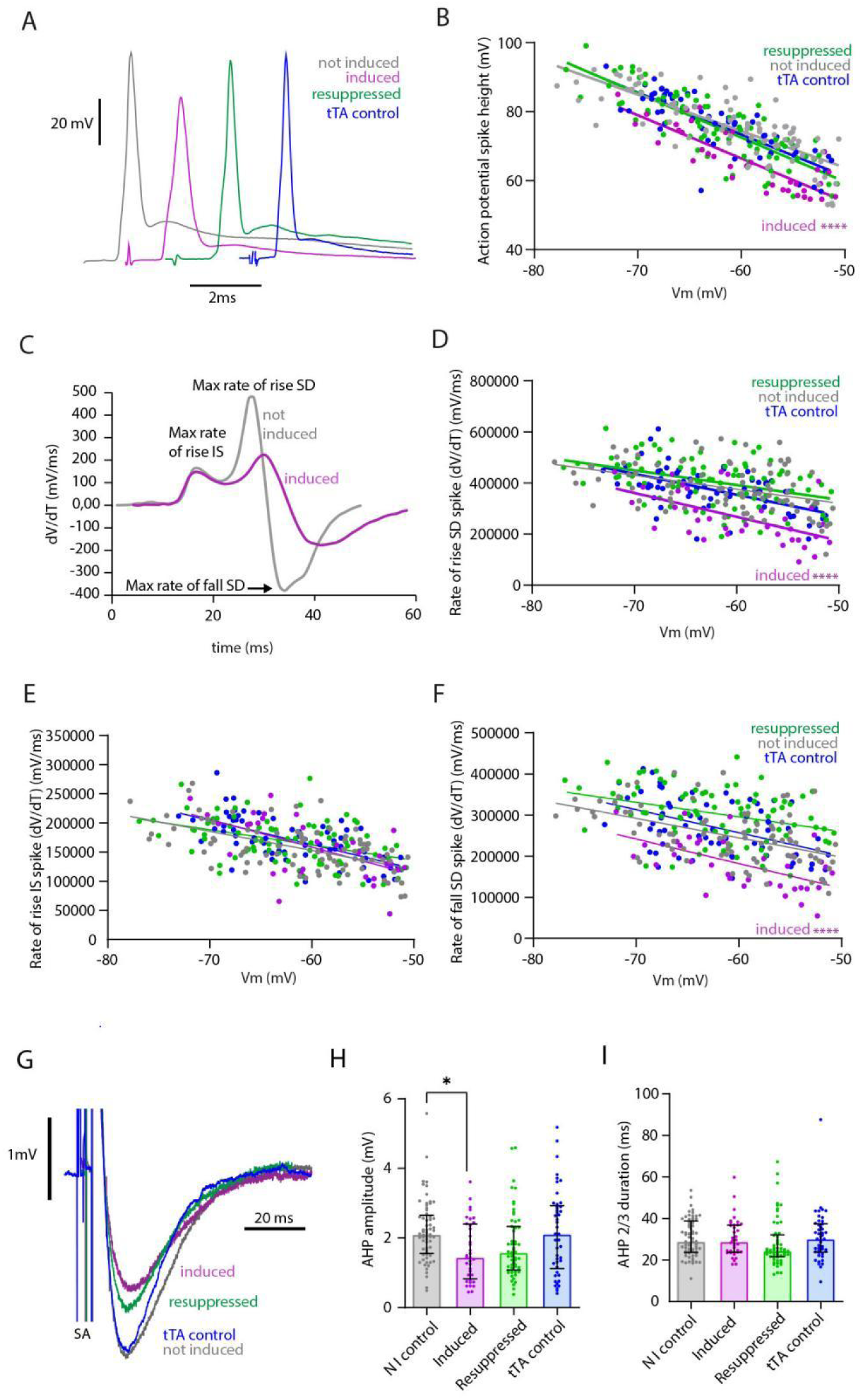
TDP-43 pathology drives decreases in action potential height and AHP amplitudes. (**A**) Representative examples of averages of antidromic action potentials recorded from motoneurones from Not-induced (grey), Induced (magenta), Resuppressed (green) and tTA control (blue). (**B**) X-Y plots showing linear regression for antidromic action potential amplitude against membrane potential. Linear regression for all 4 slope P<0.0001 (Not-induced R^2^ = 0.604, Induced R^2^ = 0.713, Resuppressed R^2^ = 0.685, tTA control R^2^ = 0.675). Not-induced: n = 87 cells (6 mice, 3 female, 3 male), Induced: n= 36 (9 mice, 4 female, 5 male), Resuppressed: n= 86 cells (6 mice, 3 female, 3 male) and tTA control: n= 63 cells (5 mice, 3 female, 2 male). The angle of the slopes were not significantly different between groups but the elevation was (P<0.0001) showing that the amplitudes of the action potential from the soma was significantly lower in the induced mice than both not-induced bigenic mice (P<0.0001) and tTA controls but returned to normal values upon resuppression. (**C**) Representative examples of the first derivatives of an antidromic action potential from a not-induced mouse (grey) and an induced mouse (magenta). On this is indicated the maximum rate of rise for both the initial segment (IS) and soma-dendritic (SD) portions of the spike and the maximum rate of fall for the somatic spike. (**D**) X-Y plots showing linear regression for the maximum rate of rise of the SD portion of the spike by membrane potential. Linear regression for induced group P=0.003, the rest all P<0.0001 (Not-induced R^2^ = 0.167, Induced R^2^ = 0.322, Resuppressed R^2^ = 0.1631, tTA control R^2^ = 0.307). The numbers of cells/mice were the same as for B. The angle of the slopes were not significantly different between groups but the elevation was (P<0.0001) showing that the decreased spike height shown in B is due to a slower rate of rise of the somatic portion of the antidromic spike which returns to normal values upon resuppression. (**E**) X-Y plots showing linear regression for the maximum rate of rise of the IS portion of the spike by membrane potential. Linear regression for induced group P=0.002, the rest all P<0.0001 (Not-induced R^2^ = 0.334, Induced R^2^ = 0.347, Resuppressed R^2^ = 0.189, tTA control R^2^ = 0.3403). The numbers of cells/mice were the same as for B. Neither the angle nor the elevation of the slopes were significantly different between groups (P=0.172 and P= 0.188 respectively). (**F**) X-Y plots showing linear regression for the maximum rate of fall for the SD portion of the spike by membrane potential. Linear regression for induced group P=0.003, the rest all P<0.0001 (Not-induced R^2^ = 0.252, Induced R^2^ = 0.337, Resuppressed R^2^ = 0.124, tTA control R^2^ = 0.223). The numbers of cells/mice were the same as for B. The angle of the slopes were not significantly different between groups (P= 0.493) but the elevations were (P<0.0001) confirming that the rate of fall of the somatic spikes were significantly slower in induced mice but returned to normal values upon resuppression. (**G**) Representative examples of AHPs recorded from motoneurones from a not-induced (grey), induced (magenta), resuppressed (green) and tTA control (blue) mice using an intracellular current pulse to evoke a single spike. Traces shown are averages of at least 10 action potentials. (**H**) Scatter dot plot showing AHP amplitudes for the four different groups. Plot shows medians and interquartile ranges and each dot shows a single neurone. Medians (and IQR), Not-induced: 2.10 mV (1.10), Induced: 1.44 mV (1.57), Resuppressed: 1.58 mV (1.25), tTA control: 2.11 mV (1.8). Kruskal Wallis, P =0.0049, Not-induced: n = 65 cells (6 mice, 3 female, 3 male), Induced: n= 36 (8 mice, 4 female, 4 male), Resuppressed: n= 55 cells (5 mice, 2 female, 3 male) and tTA control: n=46 cells (5 mice, 3 female, 2 male). Dunn’s Multiple comparisons tests: Not-induced vs. Induced P= 0.0139. Other pairwise comparisons were not significant. (**I**) Scatter dot plot showing the AHP duration measured at 2/3 amplitude for the four different groups. Plot shows medians and interquartile ranges and each dot shows a single neurone. Medians (and IQR), Not-induced: 28.9 ms (15.2), Induced: 28.8 ms (13.1), Resuppressed: 24.4 ms (10.5), tTA control: 30.1 ms (13.6). Kruskal Wallis, P =0.0906, Not-induced: n = 65 cells (6 mice, 3 female, 3 male), Induced: n= 36 (8 mice, 4 female, 4 male), Resuppressed: n= 58 cells (5 mice, 2 female, 3 male) and tTA control: n=46 cells (5 mice, 3 female, 2 male). *P < 0.05, **P < 0.01, ***P < 0.001, ****P < 0.0001

As potassium channel dysfunction has repeatedly been implicated in ALS, we also measured the maximal rate of fall of the SD spike which was also significantly lower in the induced mice than both control groups and again this was reversed upon transgene resuppression (**Fig. 5F**). Additionally, the amplitude and duration of the post-spike after-hyperpolarisation (AHP) was measured (see examples in **Fig. 5G**). The duration was unchanged (**Fig. 5I**), but the amplitude was found to be significantly reduced in the induced mice (**Fig. 5H**), although this feature was not completely reversed by resuppression (**Fig. 5H**).

## Discussion

Our results show that TDP-43 pathology is sufficient to drive an extreme hyperexcitability of spinal motoneurones in vivo, consistent with the observed excitability seen in patients. The excitability changes seen including reductions in rheobase, increases in gain, reductions in AHPs and increased persistent inward current are the same as have been observed in symptomatic SOD1 models of the disease ^10,11,15^. The anatomical changes including the elongation and constriction of the axon initial segment have also been reported in the symptomatic stages of the disease in SOD1 models ^12^. This suggests that increases in motoneurone excitability is a key feature across both familial and sporadic ALS and that the anatomical changes driving these changes in intrinsic hyper-excitability may also be common across the disease, despite the initial factors driving this in the first place. Therefore, we conclude that TDP-43 may be the common denominator explaining motoneurone hyper-excitability across almost all ALS patients.

Our results are consistent with and perfectly complement the recent work showing axon initial segment elongation and increased excitability of human iPSC cells harboring TDP-43 mutations found in this disease^21^. We here confirm that such changes are occurring with disease progression *in vivo* in the adult and are not due to a differential vulnerability of the mutant cells to in vitro conditions. Conversely, the results of Harley et al. confirm that what we are observing in our mouse models is most likely occurring in humans with the disease. Taken together, the results of the two studies definitively implicate TDP-43 as an upstream driver of axon initial segment plasticity in ALS, explaining the excitability changes seen in lower motoneurones in individuals with ALS including spasticity and fasciculations.

We also show changes in the diameters of proximal axons with both swelling at the axon hillocks and a distal constriction of the axon initial segments. Proximal swellings have also been shown on ventral horn neurones in post mortem tissue in ALS ^26^. Electron microscopy revealed these swellings to mainly consist of accumulations of 10-nm neurofilaments, but later studies from the same group also showed an accumulation of mitochondria and lysosomes ^27^. More recent work using iPSC cells from patients with C9orf72 and SOD1 mutations confirm an accumulation of phosphorylated heavy/medium neurofilament chains in axon hillocks of motoneurones in both mutations ^28^. Whilst TDP pathology is not generally associated with SOD1 ALS, it is a key feature of C9orf72 ALS. This accumulation may be due to deficits in retrograde transport, which is well documented in ALS ^29^.

TDP-43 is known to play an important role in axonal transport which has been shown to be impaired in mice carrying ALS-associated TDP-43 mutations ^30^. In our tracing studies we frequently observed the strongest fluorescence signals from the proximal axonal bulges and in many cases the tracer failed to enter the soma consistent with deficits in axonal transport. The breakdown of the proximal border of the axon initial segment has not been previously observed in SOD1 mouse models, although this has not been examined closer to end-stage. Intriguingly, this phenomenon has been observed with expression of pseudo-acetylated Tau species in models of Alzheimers Disease ^31^ suggesting that disruptions of axon initial segments may be a common feature across different neurodegenerative diseases.

By contrast the distal axon initial segments in our induced mice were of significantly smaller diameter which could be predicted to increase the excitability ^32^. This feature does not appear to return to normal values, although this constriction was also seen in the tTA only controls. One potential explanation for this could be that both the resuppressed and the tTA only controls were around 7 weeks older than the not-induced and four-week induced groups as they all started at the same ages.

We have previously seen that the diameter of axon initial segments decreases on spinal motoneurones in C57BL/6 mice between 22 days of age and ∼170 days of age by around 12%, with no changes in axon initial segment length ^13^. Therefore, it is plausible that the 13% reduction in diameter seen in the tTA mice compared to the younger not-induced controls reflects the normal maturation process and, as such, the similarity with measurements in the resuppressed mice may, in fact, reflect a return to normal of this feature also. Alternatively, it could also partly reflect a permanent loss of large diameter motor axons with functional recovery coming from re-innervation of thinner diameter axons^33^.

Our work was restricted to the spinal motoneurones, however increased excitability has also been observed in upper motoneurones in the same TDP-43 ΔNLS mouse but with the transgene driven by a CAMKIIα promoter ^34^. If similar changes are also occurring in axon initial segments of corticospinal motoneurones this could also explain their hyper-excitability and indeed, also explain lower thresholds for activation of the motor cortex observed with transcranial magnetic stimulation in humans with ALS ^2,35^.

Our observed changes are unlikely to be simply due to selective survival as many values obtained for the different measures lie far outside of the normal ranges seen in control mice. Furthermore, cell loss has been estimated to be minimal even at this disease stage (post induction time point) in this mouse model ^23^. This does not preclude a dying back of axons. Our tracing ensured that at the time of tracing the motoneurones were still functionally connected at the neuromuscular junction. This is important as both axonal injury or a loss of functional connectivity at the neuromuscular junction has been shown to drive changes to axon initial segments of spinal motoneurones ^25,36^. Thus, the changes we have observed are a result of the TDP-43 pathology rather than a consequence of the dying back process. Unlike the IPSC experiments of^21^ our axon initial segment elongation was not followed by a later axon initial segment shortening as the mice were already close to humane endpoint. This suggests that the shortening observed in the IPSC experiments may either reflect a stage beyond our humane endpoint, or specifically sick neurons that would have denervated the neuromuscular junction in vivo.

Another strong indication that our results are unlikely to be due to selective survival is perhaps one of the most striking observations in our experiments-that almost all of the changes were reversible upon resuppression of the transgene and the restoration of normal nuclear TDP-43. This observation is particularly relevant as, while a wide range of drugs have had some effect on disease survival in ALS mouse models, no experimental drug to date has been able to reverse symptoms. Drugs targeting excitability, such as riluzole, also have only limited effect in patients. One reason may be that their effects cannot differentiate between pathological excitability changes and homeostatic excitability changes crucial to maintain motor function as cortical motoneurones also die back. Our results suggest that targeting TDP-43 may represent a safer and more effective way to control excitability within healthy but still functional levels in this disease.

## Materials and methods

### Mice

Permission to conduct the following animal research was granted by the Danish Animal Experiments Inspectorate (authorization no. 2018-15-0201-01426), and all treatments conformed to the national and European laws on the welfare and protection of animals used for experimentation - i.e., the Danish Animal Welfare Act (2013) and the EU directive 2010/63/EU. Two monogenic mouse lines were used for breeding in our facilities, originally from The Jackson Laboratory. Mice constitutively expressing the tetracycline-controlled transactivator (tTA) transgene downstream of the Neurofilament Heavy Chain (NEFH) promoter (B6;C3-Tg(NEFH-tTA)8Vle/J, stock no. 025397) were crossed with mice carrying an inserted construct of human TDP-43 ΔNLS downstream of the Tet operator regulatory element (B6;C3-Tg(tetO-hTDP-43-ΔNLS)4Vle/J, stock no. 014650), and experiments were performed on the bigenic products of this cross (NEFH-tTA/tetO-hTDP-43-ΔNLS, also termed rNLS8). For these experiments, not-induced bigenic mice and monogenic (NEFH-tTA) littermates were used as controls.

All mice were fed a Doxycycline diet up to 7 weeks of age, at which point roughly 1/3 of the bigenics were maintained on this diet throughout the experiment as a bigenic suppressed (“not-induced”) control, while the remaining 2/3 of the bigenics and all monogenic tTA only controls were moved to regular chow. This led to the induction of pathological TDP-43 expression and the development of an ALS phenotype only in the bigenic animals. After four weeks of being fed normal chow, 1/2 of the bigenic ALS animals were switched back to Dox-containing chow (for resuppression of the TDP-43 pathology) together with the tTA monogenic controls, while the remaining 1/2 of the bigenic ALS animals were sacrificed at this symptomatic stage (i.e. 4 weeks post induction) together with the bigenic controls continuously kept on Dox-diet. All the mice that were re-introduced to Dox were sacrificed 6-8 weeks later.

Mice were genotyped using a standard PCR assay described in the genotyping protocols provided by the Jackson Laboratory for the Tg(NEFH-tTA) and Tg(tetO-hTDP-43-ΔNLS) constructs. Four transgene-specific primers were used for PCR amplification to enable the final detection of these sequences on electrophoretic gels; 5’-CTC GCG CAC CTG CTG AAT-3’ (Tg(NEFH-tTA) forward), 5’-CAG TAC AGG GTA GGC TGC TC-3’(Tg(NEFH-tTA) reverse), 5’-TTG CGT GAC TCT TTA GTA TTG GTT TGA TGA-3’ (Tg(tetO-hTDP-43-ΔNLS) forward) and 5’-CTC ATC CAT TGC TGC TGC G-3’ (Tg(tetO-hTDP-43-ΔNLS) reverse).

### Behavioural analysis

In the two-minute suspension test mice were suspended by their tails to identify those exhibiting an overt motor phenotype characterized by trembling and hindlimb clasping. To measure grip strength and endurance mice were placed on a wire cage lid, which was then inverted approximately 30 cm above a box filled with bedding. The endurance time that the mice could hold onto the lid before falling was measured (up to a maximum of 2 minutes), and the average of three trials was recorded weekly from one week pre induction to human endpoint or after 6-8 weeks of transgene resuppression.

### Electrophysiological experiments

These were performed as previously described ^37^. Mice were anaesthetized with a mixture consisting of Hypnorm (fentanyl-citrate 0.315 mg/mL and fluanisone 10 mg/mL), Midazolam (5 mg/mL) diluted 1:1:2 in distilled water. The body temperature was maintained throughout at 36-37°C using rectal probe controlling warming devices (a heating pad under the mouse and an infrared lamp above the mouse). IP catheters were implanted for drug delivery including the anaesthesia and the skeletal muscle relaxant Pavulon (Pancuronium bromide, diluted 1:10 with saline, given at one-hour intervals upon initiation of mechanical ventilation). A tracheal cannula was inserted for subsequent mechanical ventilation. To monitor the animal, the cardiac rhythm was recorded using electrocardiographic (ECG) traces and the partial pressure of CO2 in the exhaled air was monitored throughout the recordings.

A dorsal vertebral laminectomy (T12-L1 level) allowed access to the L3-L4 spinal segments. The sciatic nerve was dissected and stimulated using custom hook electrodes to activate action potentials in both the sensory afferent fibers and the motor fibers. This allowed for antidromic identification of spinal motoneurones impaled with a glass microelectrode filled with 2M potassium acetate solution (resistance: approx. 25 MΩ). To measure the intrinsic excitability of the impaled motoneurones, current was injected through the microelectrode in current clamp mode to mimic synaptic input, and the resulting voltage response was measured.

The intracellular signal was first amplified 10x by a headstage connected to an AxoClamp-2B amplifier using either Bridge or Discontinuous Current Clamp (DCC) modes depending on the parameter being measured. Stronger signal amplification and data filtration was achieved using Neurolog amplifiers and filters (Digitimer, UK), while analog-to-digital translation was accomplished with a CED 1401 converter (Cambridge Electronic Design, UK). Lastly, the Spike2 software (version 7.10) was used for data acquisition and analysis (Cambridge Electronic Design, UK).

Antidromic (back-fired) action potentials were identified and recorded in Bridge mode. Ten or more consecutive spikes were averaged for each impaled motoneurone, and the amplitude and baseline potential (Vm) of this average were measured in Spike2 (7.10) before it was exported to Microsoft Excel. As an inclusion criterion, it was ensured that each spike was overshooting and had a baseline potential ˂ -50 mV, confirmed using the extracellular reference potential upon electrode withdrawal from the neuron. The first derivatives of the averaged waveforms (using Microsoft Excel) were used to distinguish the initial segment (IS) and somatodendritic (SD) spike components as distinct peaks, from which the maximum rate of rise of each as well as the IS-SD latency – i.e. the time interval separating the two – could be acquired. In addition, the maximum rate of fall was determined by identifying the lowest point or absolute minimum on the derivative.

For measurements of the afterhyperpolarization (AHP), spikes were evoked by intracellular injection of brief (i.e. 1 millisecond long) positive current pulses and recorded in Bridge mode to avoid any synaptic contamination. For these recordings, the Vm was adjusted and maintained around -70 mV by constant current injection. The amplitudes were measured from averaged AHPs (using spike2) and calculated as the difference between the baseline Vm and the minimum (i.e. most negative) Vm reached in this refractory period. Next, AHP duration was calculated as the time interval between the point of transition at the baseline and the point at which the voltage went back to two-thirds of the AHP amplitude during its decay (as the exact return can be difficult to determine).

The input resistance was measured for each neurone as the voltage drop in response to an intracellular injection of a 50 milliseconds long negative current pulse with an intensity of -3 nA (using 5 kHz DCC mode) to trigger a small membrane hyperpolarization. Ten or more of these responses were averaged, and the voltage change was determined from the average in Spike2 (7.10) to determine the input resistance. In cases where a sag was observed at the end of a triggered voltage drop, the absolute and relative (% of peak hyperpolarization) size of this was also estimated to uncover a potential dysfunction of hyperpolarization-activated cyclic nucleotide-gated (HCN) channels.

Repetitive spike discharges were triggered by somatic injection of 3 and 8 kHz triangular current pulses and recorded in DCC mode. As a first step, the rheobase and the spiking threshold were obtained from these recordings for each penetrated motoneuron. Next, the instantaneous firing frequency was plotted in Spike2 (7.10) and exported to Excel to identify the primary and secondary ranges of firing based on the sudden acceleration of the firing frequency seen in the otherwise linear current-frequency (I-f) relationships. The linear trendline equations were obtained in Excel to determine the input-output gains of the motoneurones.

### Anatomical tracing experiments

Under isoflurane anaesthesia (1.5-2%) a small incision was made at the Achilles tendon and the tracer Fast Blue (FB, Polysciences, 1.5% in 3 µL PBS) was injected into the soleus muscle and Alexa Fluor 488-conjugated Cholera Toxin subunit B (CTB 488, Thermo Fisher Scientific, 0.05% in 5 µL PBS) into the gastrocnemius muscle. These represent muscles with greater proportions of slow or faster motor units respectively.

Pain management was obtained peri-operatively with buprenorphine (injected subcutaneously, 0.1 mg/kg), and post-operatively with buprenorphine delivered orally (mixed in Nutella, 5 mg/g /12 hours) for three days after the surgery. A single dose of the antibiotic tribrissen (trimethoprim-sulfadiazine) was given peri-operatively (dose: 0.1 mL of stock solution: 40 mg trimethoprim/200 mg sulfadiazine/mL diluted 1:10).

### Immunohistochemistry

Immediately after the electrophysiological experiments, the mice were given an euthanizing dose of sodium pentobarbital (I.P, 120 mg/kg). Mice were then transcardially perfused with saline followed by 4% paraformaldehyde in phosphate buffer, (pH 7.4).The spinal cord was post-fixed for 2 hours in the same fixative mixed with 15% sucrose, then stored in 30% sucrose dissolved in phosphate buffered saline. The spinal cord was then horizontally sliced into 50 µm thick sections on a freezing microtome (Thermo Scientific, HM450). Immunohistochemistry was performed on free-floating ventral sections blocked in 5% Donkey Serum. To label axon initial segments, sections were incubated with a primary antibody targeting Ankyrin G overnight (rabbit anti-AnkG, 1:500, H215, Santa Cruz Cat no: sc28561) followed by 2 hours incubation in the secondary antibody (Alexa FluorTM 568 donkey anti-rabbit, 1:1000, Thermo Scientific Cat no: A-11055) diluted in PBST. Sections were mounted on glass slides and coverslipped using Prolong Gold Antifade mounting medium. In some experiments double labelling was also performed to label C-boutons for another project ^38^.

### Imaging and measuring the axon initial segments

3-dimensional measurements were made from Z-stacks. These were captured by confocal microscopy (ZEISS LSM700) and using Zen software (black edition, Zeiss). For traced motoneurones the length and distal diameter of the axon initial segment was measured as well as the distance between the cell body and the proximal border of the axon initial segment. The 2D soma size was approximated by measuring the longest and shortest diameters in a 2-dimensional plane and calculating the area of an ellipse from these. Unlike in normal motoneurones, where the proximal and distal borders are easy to manually identify, in induced animals we frequently observed examples where the proximal borders of the axon initial segment were dissipated. Fluorescence intensity profiles were therefore used for this purpose (still using ZEN software). Representative images were prepared for publication using FIJI (ImageJ, NIH), Images for the figures were prepared using Fiji (Image J, NIH) and Adobe Illustrator. Brightness, contrast, gaussian blur and background subtraction were adjusted for image presentation using Image J and this was performed uniformly across the entire image.

### Statistics

D’Agostino Omnibus tests were conducted in GraphPad prism (version 9). Data that were normally distributed were analysed by means of parametric ANOVAS (followed by post hoc Tukey tests), while non-normally distributed data were analysed using Kruskal-Wallis tests (followed by post hoc Dunns Multiple comparison tests). Between animal variation even within a group is expected due to the variation of motor unit subtypes (fast vs. slow) and the inability to equally sample from identical proportions of these with in vivo intracellular recording. Thus, analyses are performed by cell rather than by animal.

Linear associations between some of the measured variables (for intrinsic properties influenced by the baseline potential) were analysed using simple linear regression to reveal any potential differences in the angle of the slopes or their elevations. The critical threshold for statistical significance was set to 0.05 (5%), and the degrees of significance are denoted by asterisks for four different probability levels in the visual representations of the data (see results); *p ≤ 0.05, **p ≤ 0.01, ***p ≤ 0.001 and ****p ≤ 0.0001

Blinding was attempted during analyses but in reality due the extreme excitability levels and abnormal accumulation of the tracer in axon hillocks in the induced TDP-43 ΔNLS mice this was not possible. The necessary sample sizes with respect to animals was selected based on previous analyses of similar parameters in other ALS mouse models using similar techniques based on the average number of cells we can normally penetrate per mouse.

## Data availability

The final data from the accepted manuscript will be made available in an online repository at the University of Copenhagen. Additional data, material, and protocols are provided upon reasonable request to the corresponding author.

## Acknowledgments

All imaging was performed at the Core Facility for Integrated Microscopy, Faculty of Health and Medical Sciences, University of Copenhagen. We thank Sif Møbjerg Jørgensen for assistance with the initial training of animals.

## Funding

ANB received a scholar stipend from the Independent Research Fund Denmark. ZZ received a PhD scholarship from the China Scholarship Council. This research was funded by The Lundbeck Foundation (R370-2021-1109).

## Competing interests

The authors report no competing interests.

## Notes

### Competing Interest Statement

The authors have declared no competing interest.

### Summary of Updates

We added a special symbol delta missing in the title due to a typo error.

## References

1. Christensen, P.B., Nielsen, J.F. & Sinkjaer, T. Quantification of hyperreflexia in amyotrophic lateral sclerosis (ALS) by the soleus stretch reflex. Amyotroph Lateral Scler Other Motor Neuron Disord 4, 106–111 (2003).

2. Geevasinga, N., Van den Bos, M., Menon, P. & Vucic, S. Utility of Transcranial Magnetic Simulation in Studying Upper Motor Neuron Dysfunction in Amyotrophic Lateral Sclerosis. Brain Sci 11(2021).

3. Kanai, K., et al. Altered axonal excitability properties in amyotrophic lateral sclerosis: impaired potassium channel function related to disease stage. Brain 129, 953–962 (2006).

4. Soliven, B. & Maselli, R.A. Single motor unit H-reflex in motor neuron disorders. Muscle Nerve 15, 656–660 (1992).

5. Tankisi, H., et al. Three different short-interval intracortical inhibition methods in early diagnosis of amyotrophic lateral sclerosis. Amyotroph Lateral Scler Frontotemporal Degener 24, 139–147 (2023).

6. Vucic, S. & Kiernan, M.C. Axonal excitability properties in amyotrophic lateral sclerosis. Clin Neurophysiol 117, 1458–1466 (2006).

7. Vucic, S., Nicholson, G.A. & Kiernan, M.C. Cortical hyperexcitability may precede the onset of familial amyotrophic lateral sclerosis. Brain 131, 1540–1550 (2008).

8. Zanette, G., et al. Different mechanisms contribute to motor cortex hyperexcitability in amyotrophic lateral sclerosis. Clin Neurophysiol 113, 1688–1697 (2002).

9. Zanette, G., et al. Changes in motor cortex inhibition over time in patients with amyotrophic lateral sclerosis. J Neurol 249, 1723–1728 (2002).

10. Jensen, D.B., Kadlecova, M., Allodi, I. & Meehan, C.F. Spinal motoneurones are intrinsically more responsive in the adult G93A SOD1 mouse model of amyotrophic lateral sclerosis. J Physiol 598, 4385–4403 (2020).

11. Jensen, D.B., Kadlecova, M., Allodi, I. & Meehan, C.F. Response to Letter to Editor on the article Jensen DB, Kadlecova M, Allodi I, Meehan CF (2020). J Physiol 599, 4233–4236 (2021).

12. Jørgensen, H.S., et al. Increased Axon Initial Segment Length Results in Increased Na(+) Currents in Spinal Motoneurones at Symptom Onset in the G127X SOD1 Mouse Model of Amyotrophic Lateral Sclerosis. Neuroscience 468, 247–264 (2021).

13. Bonnevie, V.S., et al. Shorter axon initial segments do not cause repetitive firing impairments in the adult presymptomatic G127X SOD-1 Amyotrophic Lateral Sclerosis mouse. Sci Rep 10, 1280 (2020).

14. Delestrée, N., et al. Adult spinal motoneurones are not hyperexcitable in a mouse model of inherited amyotrophic lateral sclerosis. J Physiol 592, 1687–1703 (2014).

15. Huh, S., Heckman, C.J. & Manuel, M. Time Course of Alterations in Adult Spinal Motoneuron Properties in the SOD1(G93A) Mouse Model of ALS. eNeuro 8(2021).

16. Meehan, C.F., et al. Intrinsic properties of lumbar motor neurones in the adult G127insTGGG superoxide dismutase-1 mutant mouse in vivo: evidence for increased persistent inward currents. Acta Physiol (Oxf) 200, 361–376 (2010).

17. Dukkipati, S.S., Garrett, T.L. & Elbasiouny, S.M. The vulnerability of spinal motoneurons and soma size plasticity in a mouse model of amyotrophic lateral sclerosis. J Physiol 596, 1723–1745 (2018).

18. Suzuki, N., Nishiyama, A., Warita, H. & Aoki, M. Genetics of amyotrophic lateral sclerosis: seeking therapeutic targets in the era of gene therapy. J Hum Genet 68, 131–152 (2023).

19. Neumann, M., et al. Ubiquitinated TDP-43 in frontotemporal lobar degeneration and amyotrophic lateral sclerosis. Science 314, 130–133 (2006).

20. Suk, T.R. & Rousseaux, M.W.C. The role of TDP-43 mislocalization in amyotrophic lateral sclerosis. Mol Neurodegener 15, 45 (2020).

21. Harley, P., et al. Aberrant axon initial segment plasticity and intrinsic excitability of ALS hiPSC motor neurons. Cell Rep 42, 113509 (2023).

22. Weskamp, K., et al. Shortened TDP43 isoforms upregulated by neuronal hyperactivity drive TDP43 pathology in ALS. J Clin Invest 130, 1139–1155 (2020).

23. Walker, A.K., et al. Functional recovery in new mouse models of ALS/FTLD after clearance of pathological cytoplasmic TDP-43. Acta Neuropathol 130, 643–660 (2015).

24. Evans, M.D., Dumitrescu, A.S., Kruijssen, D.L.H., Taylor, S.E. & Grubb, M.S. Rapid Modulation of Axon Initial Segment Length Influences Repetitive Spike Firing. Cell Rep 13, 1233–1245 (2015).

25. Jensen, D.B., Klingenberg, S., Dimintiyanova, K.P., Wienecke, J. & Meehan, C.F. Intramuscular Botulinum toxin A injections induce central changes to axon initial segments and cholinergic boutons on spinal motoneurones in rats. Sci Rep 10, 893 (2020).

26. Sasaki, S. & Maruyama, S. Increase in diameter of the axonal initial segment is an early change in amyotrophic lateral sclerosis. J Neurol Sci 110, 114–120 (1992).

27. Sasaki, S. & Iwata, M. Impairment of fast axonal transport in the proximal axons of anterior horn neurons in amyotrophic lateral sclerosis. Neurology 47, 535–540 (1996).

28. Lefebvre-Omar, C., et al. Neurofilament accumulations in amyotrophic lateral sclerosis patients’ motor neurons impair axonal initial segment integrity. Cell Mol Life Sci 80, 150 (2023).

29. Nagano, S. & Araki, T. Axonal Transport and Local Translation of mRNA in Amyotrophic Lateral Sclerosis. in Amyotrophic Lateral Sclerosis (ed. Araki, T.) (Exon Publications, Brisbane (AU), 2021).

30. Sleigh, J.N., et al. Mice Carrying ALS Mutant TDP-43, but Not Mutant FUS, Display In Vivo Defects in Axonal Transport of Signaling Endosomes. Cell Rep 30, 3655–3662.e3652 (2020).

31. Sohn, P.D., et al. Pathogenic Tau Impairs Axon Initial Segment Plasticity and Excitability Homeostasis. Neuron 104, 458–470.e455 (2019).

32. Halter, J.A. & Clark, J.W., Jr. The influence of nodal constriction on conduction velocity in myelinated nerve fibers. Neuroreport 4, 89–92 (1993).

33. Hur, S.K., et al. Slow motor neurons resist pathological TDP-43 and mediate motor recovery in the rNLS8 model of amyotrophic lateral sclerosis. Acta Neuropathol Commun 10, 75 (2022).

34. Dyer, M.S., et al. Mislocalisation of TDP-43 to the cytoplasm causes cortical hyperexcitability and reduced excitatory neurotransmission in the motor cortex. J Neurochem 157, 1300–1315 (2021).

35. Dharmadasa, T. Cortical Excitability across the ALS Clinical Motor Phenotypes. Brain Sci 11(2021).

36. Kiryu-Seo, S., et al. Impaired disassembly of the axon initial segment restricts mitochondrial entry into damaged axons. Embo j 41, e110486 (2022).

37. Meehan, C.F., Sukiasyan, N., Zhang, M., Nielsen, J.B. & Hultborn, H. Intrinsic properties of mouse lumbar motoneurons revealed by intracellular recording in vivo. J Neurophysiol 103, 2599–2610 (2010).

38. Bak, A.N., et al. Cytoplasmic TDP-43 accumulation drives changes in C-bouton number and size in a mouse model of sporadic Amyotrophic Lateral Sclerosis. Mol Cell Neurosci 125, 103840 (2023).

