## Supplement Figure 1 for "TDP-43 pathology is sufficient to drive axon initial segment plasticity and hyperexcitability of spinal motoneurones in vivo in the TDP43-ΔNLS model of Amyotrophic Lateral Sclerosis"

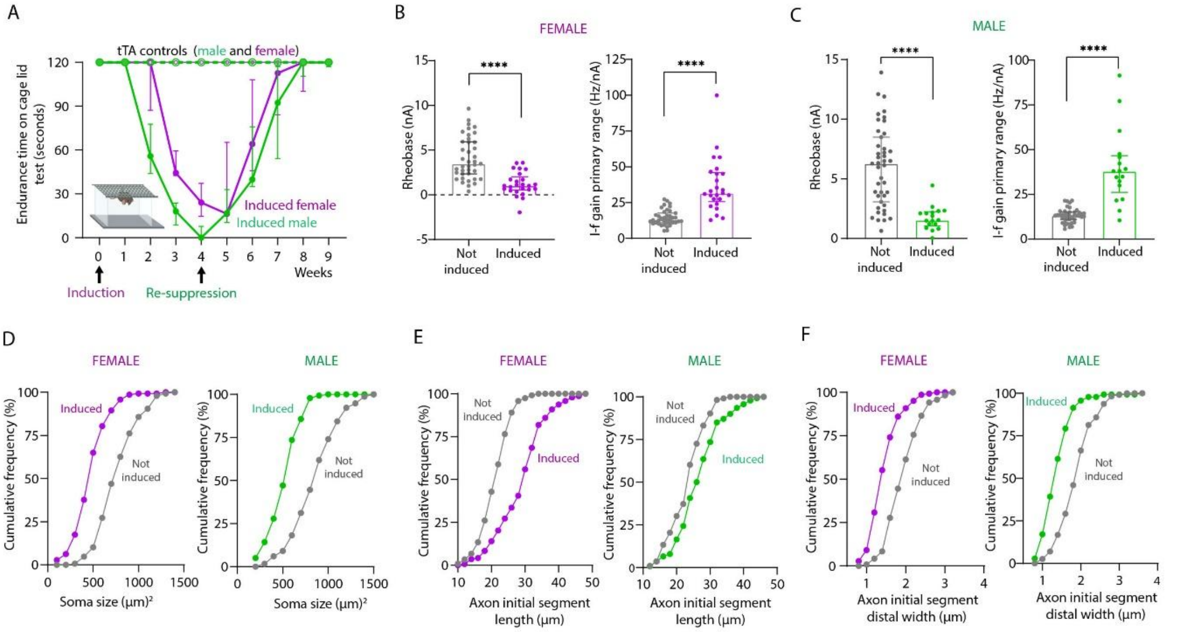


**Supplement Figure 1: Behavioral, structural and electrophysiological characteristics by sex.** (**A**) Graph showing the endurance time on the cage grid test for induced TDP-43 ΔNLS mice and tTA only controls going through the same doxycycline on, off and on cycle. From this it can be seen that the male induced mice (green) show a reduced endurance time earlier after induction than induced female mice (magenta). In fact, 2 male mice had to be resuppressed 3 days earlier than the others to prevent them reaching humane endpoint. Both sex recover this motor unction after resuppression n=9 tTA mice (5 female, 4 male) and n=12 induced ΔNLS mice (4 female, 8 male). (**B**) Scatter dot plots showing recruitment currents (rheobase) and the current-frequency (I-f) gains measured in the primary range from not-induced (grey) and induced (magenta) groups in female mice. This shows that the rheobase decreased significantly and the I-f gain increase significantly in female induced mice. Female rheobase means (and SD), Not-induced: 4.11 nA (2.37), Induced: 1.22 nA (1.30), t-test, P <0.0001, Not-induced: n = 39 cells (3 mice), Induced: n= 25 (5 mice). Female I-f gain medians (and IQR), Not-induced: 13.04 Hz/nA (6.66), Induced: 31.53 Hz/nA (20.12), Mann Whitney, P <0.0001, Not-induced: n = 39 cells (3 mice), Induced: n= 25 cells (5 mice). Plot shows medians and interquartile ranges and each dot shows the rheobase of a single neurone. (**C**) Scatter dot plots showing recruitment currents (rheobase) and the current-frequency (I-f) gains measured in the primary range from not-induced (grey) and induced (magenta) groups in male mice. This shows that the rheobase also decreased significantly and the I-f gain increased significantly in male induced mice. Male rheobase medians (and IQR), Not-induced: 6.29 nA (5.43), Induced: 1.55nA (1.2), Mann Whitney, P <0.0001, Not-induced: n = 43 cells (3 mice), Induced: n= 17 cells (4 mice). Male I-f gain means (and SD), Not-induced: 13.11 Hz/nA (23.88), Induced: 40.14 Hz/nA (20.73), t-test, P <0.0001, Not-induced: n = 43 cells (3 mice), Induced: n= 17 (4 mice). Plot shows medians and interquartile ranges and each dot shows the rheobase of a single neurone. (**D**) Cumulative frequency distribution of soma size by gender for traced motoneurones (both muscles). Left panel: female groups, right panel: male groups. These show that both sexes exhibited a significant decrease in soma size after transgene induction. Females medians (and IQR), Not-induced: 773.4 $\mu m^2$ (311.4), Induced: 495.3 $\mu m^2$ (210.9), Mann Whitney, P <0.0001, Not-induced: n = 147 cells (4 mice), Induced: n= 143 cell (5 mice). Male means (and SD), Not-induced: 887.1 $\mu m^2$ (256.6), Induced: 554.4 $\mu m^2$ (171.5), t-test, P <0.0001, Not-induced: n = 231 cells (4 mice), Induced: n= 140 (3 mice). (**E**) Cumulative frequency distribution of axon initial segment length for traced motoneurones (both motor pools collapsed) by gender. Left panel: female group, right panel: male group. These show that both sexes exhibited a significant lengthening of axon initial segments after transgene induction but this was more pronounced in female mice. Female means (and SD), Not-induced: 21.81 µm (3.423), Induced: 29.53 µm (7.33), t-test, P <0.0001, Not-induced: n = 119 cells (4 mice), Induced: n= 143 cells (5 mice). Male means (and SD), Not-induced: 23.84 µm (5.164), Induced: 27.30 µm (6.92), t-test, P <0.0001, Not-induced: n = 113 cells (4 mice), Induced: n= 140 cells (3 mice). (**F**) Cumulative frequency distribution of axon initial segment distal width by gender. Left panel: female group, right panel: male group. Females medians (and IQR), Not-induced: 1.98 µm (0.65), Induced: 1.44 µm (0.5), Mann- Whitney, P <0.0001, Not-induced: n = 119 cells (4 mice), Induced: n= 143 cells (5 mice). Males medians (and IQR), Not-induced: 1.98 µm (0.58), Induced: 1.4 µm (0.52), Mann- Whitney, P <0.0001, Not-induced: n = 113 cells (4 mice), Induced: n= 140 cells (3 mice).
